# Multiscale Computational Framework to Investigate Integrin Mechanosensing and Cell Adhesion

**DOI:** 10.1101/2023.03.24.533575

**Authors:** Andre R. Montes, Gabriela Gutierrez, Adrian Buganza Tepole, Mohammad R.K. Mofrad

## Abstract

Integrin mechanosensing plays an instrumental role in cell behavior, phenotype, and fate by transmitting mechanical signals that trigger downstream molecular and cellular changes. For instance, force transfer along key amino acid residues can mediate cell adhesion. Disrupting key binding sites within *α*_5_*β*_1_ integrin’s binding partner, fibronectin (FN) diminishes adhesive strength. While past studies have shown the importance of these residues in cell adhesion, the relationship between the dynamics of these residues and how integrin distributes force across the cell surface remains less explored. Here, we present a multiscale mechanical model to investigate the mechanical coupling between integrin nanoscale dynamics and whole-cell adhesion mechanics. Our framework leverages molecular dynamics simulations to investigate residues within *α*_5_*β*_1_-FN during stretching and the finite element method to visualize the whole-cell adhesion mechanics. The forces per integrin across the cell surface of the whole-cell model were consistent with past atomic force microscopy and Förster resonance energy transfer measurements from literature. The molecular dynamics simulations also confirmed past studies that implicate two key sites within FN that maintain cell adhesion: the synergy site and RGD motif. Our study contributed to our understanding of molecular mechanisms by which these sites collaborate to mediate whole-cell integrin adhesion dynamics. Specifically, we showed how FN unfolding, residue binding/unbinding, and molecular structure contribute to *α*_5_*β*_1_-FN’s nonlinear force-extension behavior during stretching. Our computational framework could be used to explain how the dynamics of key residues influence cell differentiation or how uniquely designed protein structures could dynamically limit the spread of metastatic cells.

## I. INTRODUCTION

Cell-matrix junctions, governed in part by macromolecular structures known as focal adhesions (FAs), can alter cell phenotype, behavior, and fate via applied mechanical signals that trigger downstream molecular and cellular changes^1–9^. At the heart of FA formation is a transmembrane heterodimer known as integrin containing *α*- and *β* - subunits. Normally, nascent FAs initiate with integrin activation, where cytoplasmic proteins bind to the integrin tails and the integrin head extends to an active state with a higher affinity for ligand binding^2,10^. However, the activation of a particular integrin, *α*_5_*β*_1_ appears to follow a separate mechanism where an extended conformation may not be required to bind to its primary ligand, fibronectin (FN)^11,12^. Instead, *α*_5_*β*_1_ binds to FN before cytoplasmic proteins anchor it to the cytoskeleton and additional integrins cluster together to create a mature FA (Fig. 1A).

**FIG. 1.**
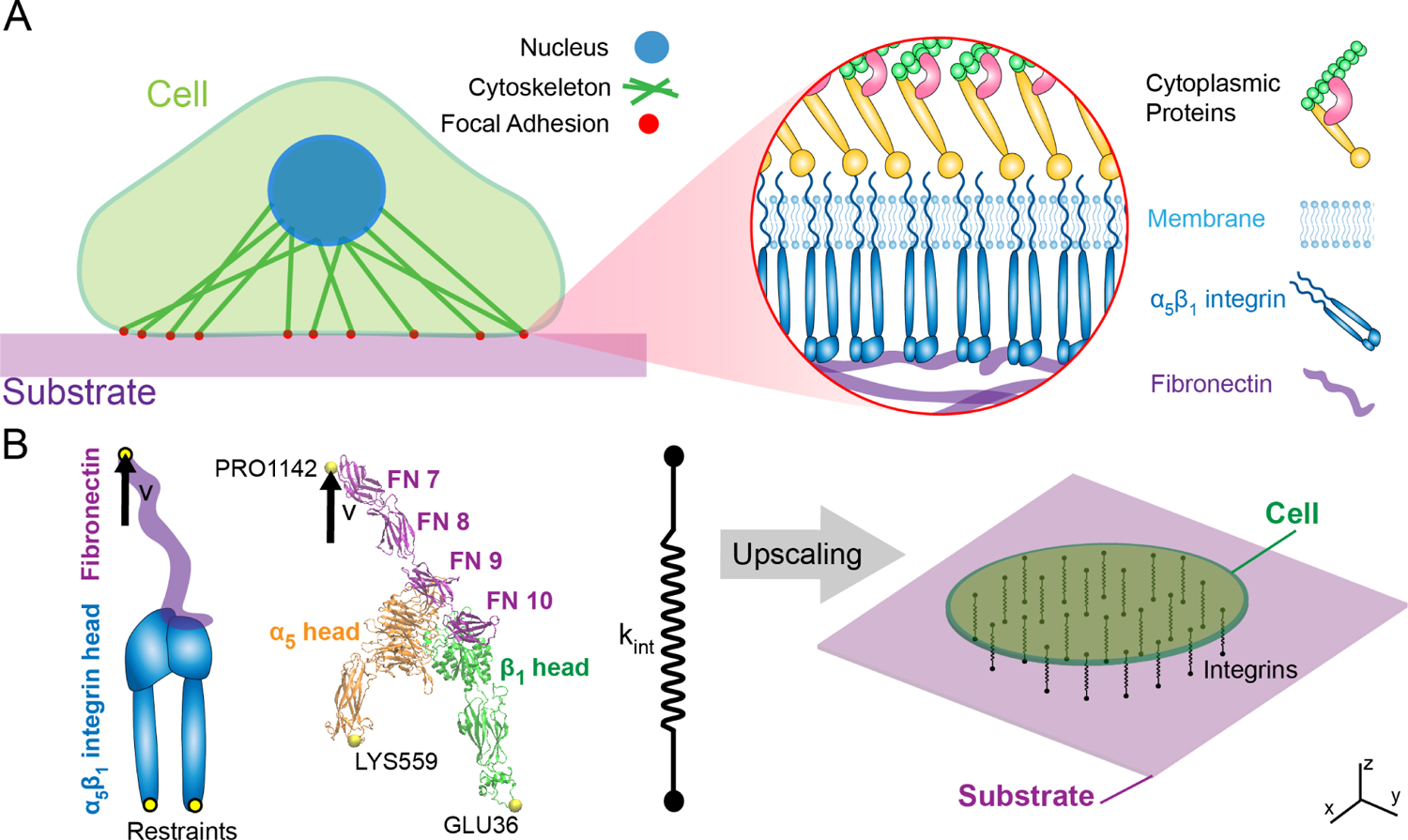
Simplified schematic of multiscale cell mechanobiology within cell adhesion mediated by *α*_5_*β*_1_ integrin (A) The cell attaches to a substrate via FAs which house multiple biomolecules including cytoplasmic proteins that anchor integrins to corresponding ligands. (B) The molecular assembly consisted of *α*_5_*β*_1_ integrin head bound to fibronectin type III fragment 7-10. For the MD simulations, restraints were placed on GLU36 and LYS559 with an applied velocity at PRO1142. The *α*_5_*β*_1_-FN’s stretching behavior was characterized by a spring that was applied to a 2D continuum model of an elastic cell on a substrate.

The connection between *α*_5_*β*_1_ integrin and FN is a main mechanosensing unit for external forces transmitting along amino acid residues that mediate cell adhesion^12^. The two principal *α*_5_*β*_1_ binding sites in FN include the 8-amino-acid-long DRVPHSRN synergy site and the RGD motif^12–14^. Upon mutation of R1374 and R1379 within the synergy site, spinning disk assays showed a reduction in cell-substrate adhesion strength; moreover, a perturbation of FN’s RGD motif inhibited adhesion altogether^15^. While the synergy site and RGD motif have been shown to play a role in cell adhesion, their nanoscale dynamics and force transduction pathway are less resolved. Elucidating how these residues maintain cell adhesion during integrin mechanosensing is important because their nanomechanics could be leveraged to control cell phenotype or motility.

Notably, *α*_5_*β*_1_’s predominant role in mediating cell adhesion lends itself to be instrumental in the progression of various pathologies. For example, imposing a fibrotic microenvironment on cells by depositing collagen-I or applying biomechanical forces to the cancer cells leads to greater *α*_5_*β*_1_ integrin-mediated proliferation^16,17^. Similarly, as a tumor’s rigidity increases, mechanosensitive *α*_5_*β*_1_ integrins are recruited and cluster together, creating larger FAs and stress fibers that promote tumor growth via a positive biochemical and biophysical feedback loop^18,19^. By understanding the link between nano and micromechanics of the cell, we could influence differentiation or mitigate the uncontrolled spread of metastatic cells through targeted protein or drug design.

Therefore, to uncover the mechanical coupling between the nanoscale dynamics of key residues in *α*_5_*β*_1_ integrin and whole-cell adhesion dynamics, we built a multiscale model. Specifically, we combined adhesion kinetics, the finite element (FE) method, and molecular dynamics (MD) to demonstrate how key residues contributed to spring-like force-extension behavior which in turn influenced the whole-cell spatial distribution of forces on integrins (Fig. 1B). The force per integrin results from our model were within those measured by past atomic force microscopy (AFM)^20^ and Förster resonance energy transfer (FRET) measurements^21^. The model indicated localization of *α*_5_*β*_1_ integrin along the cell periphery, which is consistent with cell-based studies that stain for *β*_1_ integrin and FN fragments^22^. Most importantly, the model contributed an inside look at the molecular dynamics by which the DRVPHSRN synergy site and RGD motif work together to mediate whole-cell adhesion mechanics.

## II. METHODS

### A. All-atom Steered Molecular Dynamics

The 7NWL.pdb file containing human *α*_5_*β*_1_ integrin in complex with FN and TS2/16 Fv-clasp was downloaded from the Protein Data Bank^12^. Schumacher et al. used the TS2/16 Fv-clasp to aid in the crystallization of *α*_5_*β*_1_-FN and is not naturally occurring and was therefore removed using PyMOL 2.5^23^, leaving three protein chains to be analyzed as part of the remaining complex: *α*_5_ integrin, *β*_1_ integrin, and FN type III. We refer to this complex, or system as “*α*_5_*β*_1_-FN.”

All-atom molecular dynamics (MD) simulations were run in GROMACS 2018.3^24^ with the AMBER99SB-ildn force field and periodic boundary conditions. Using the Gromacs built-in function, gmx editconf, we rotated the *α*_5_*β*_1_-FN complex 45 degrees to align the structure inside a 18nm x 45nm x 19nm box. The structure was solvated in a TIP3P water box with 0.15mM NaCl resulting in a system with 1.5 million atoms.

The energy minimization step was carried out for 15k steps utilizing the steepest gradient descent algorithm with a step size of 0.005nm. Energy over time was extracted using the gmx energy command and then plotted in Python. The structure was then equilibrated using a sequential 1ns NVT followed by a 10ns NPT simulation with H-bonds restrained. For the NVT simulation, we used Nose-Hoover temperature coupling at 310K. For the NPT simulation, Parrinello-Rahman pressure coupling at 1 bar was added. After the equilibration runs were completed, we extracted and plotted the root-mean-square deviation (RMSD), temperature, and pressure to confirm system stability.

Upon verifying system equilibration, we ran two steered MD simulations. The positions of Lysine (LYS) 559 and glutamic acid (GLU) 36 at the proximal ends of the integrin headpieces were restrained using the gmx genrestr command (Fig. 1B). Proline (PRO) 1142 at the distal end of the FN chain was pulled vertically at 1 and 10 nm/ns using a 50kJ/mol/nm spring with an umbrella potential for 25 and 3 ns, respectively. Constant force simulations were ran with vertical pulling forces of 300 and 500 pN on PRO1142. The simulations only model the *α*_5_*β*_1_ integrin headpiece and assume that the lower legs of *α*_5_*β*_1_ and cell membrane, which are omitted, fix the positions of the headpieces at the proximal end. The model also assumes a completely vertical pulling load stemming from cell and substrate displacement and ignores any shear or rotational loads. The timestep for all steered MD simulations was 2fs. The Molecular Dynamics Parameter (.mdp) files for running the energy minimization, equilibration, and steered MD can be found in the Supplementary Materials. We used the Gromacs built-in function gmx gyrate to measure the radius of gyration of the *α*_5_ and *β*_1_ integrin heads.

### B. Force Distribution Analysis

Protein structures and MD simulation trajectories were visualized in Visual Molecular Dynamics (VMD) 1.9.4a^25^. We then used the Time-Resolved Force Distribution Analysis (FDA) software package, gromacs-fda (available: https://github.com/HITS-MBM/gromacs-fda) with Gromacs 2020.4 to calculate the punctual stresses at each of the residues along the *α*_5_ and *β*_1_ integrin chains, as well as FN. The punctual stress is the sum of absolute values of scalar pairwise forces exerted on each residue. The parameter settings for the FDA can be found in the Supplementary Materials. The gromacs-fda-vmd plugin overlaid the punctual stress heatmap onto the protein renderings in VMD. Areas of interest for the FDA were the DRVPHSRN synergy site and RGD motif/loop (Fig. 3).

### C. Whole-Cell Finite Element Model

The custom finite element (FE) model represented the cell as a thin elastic disk on top of an elastic substrate. The cell surface was assumed to be a neo-Hookean^26^ constitutive material model.

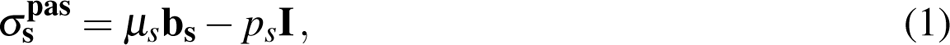

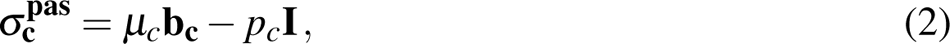

where *σ*_s_ **^pas^** and *σ***_c_ ^pas^** are the passive substrate and cell stress respectively. The shear moduli are denoted *µ_s_*, *µ_c_* (Table I). The deformation is characterized by the left Cauchy-Green tensors **b_s_**, **b_c_**. The pressures *p_s_, p_c_*are computed from boundary conditions, in this case for plane stress, ignoring 3D deformations.

**TABLE I.** Whole-cell Model Parameter Settings.

| Parameter | Variable | Setting |
| --- | --- | --- |
| Substrate modulus | $\mu_s$ | 1 MPa |
| Substrate density | $\rho_s$ | $1.0 \mu\text{g}/\mu\text{m}^3$ |
| Cell modulus | $\mu_c$ | 1 kPa |
| Cell density | $\rho_s$ | $1000 \text{ kg}/\text{m}^3$ |
| Max $\alpha_5\beta_1$-FN density | $\rho_{i_{max}}$ | $100 \mu\text{m}^{-2}$ |
| Catch-slip bond parameters | $K_a$ | $0.004 \text{ s}^{-1}$ |
| | $K_b$ | $10 \text{ s}^{-1}$ |
| | $F_a$ | $15 \text{ pN}$ |
| | $F_b$ | $15 \text{ pN}$ |

To account for cell contractility, an active stress field was applied inside the cell,

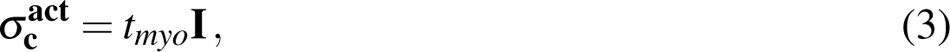

where *σt_myo_* **^act^** is the active cell stress due to the applied actin-myosin traction, *t_myo_* (Pa):

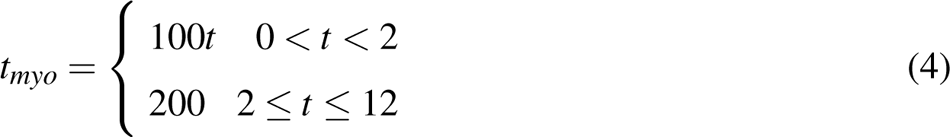

where *t* is the simulation time. We used a previously developed catch-slip bond model of adhesion to determine the number of integrin-substrate bonds per node in the FE mesh in a force dependent manner^27,28^. This model assumes that the *α*_5_*β*_1_-FN complexes behave as parallel springs that connect and disconnect to the substrate based on an association constant, *K_on_* and on a force dependent dissociation constant, *K_o f f_*, respectively.

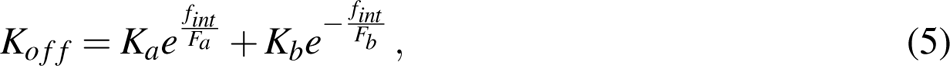

where *K_a_*, *F_a_*, *K_b_*, and *F_b_* are fitted parameters (Table I) and *f_int_* is the magnitude of the force per *α*_5_*β*_1_-FN bond. The force vector per bond, (**f_int_**), is computed via the *α*_5_*β*_1_-FN spring constant *k_int_* and the spring extension vector **u_int_**:

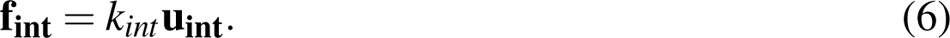

The force per node from integrin and is related to the fraction (concentration) of *α*_5_*β*_1_-FN bonds *C* with respect to the maximum density *ρ_i,max_* (Table I), the local area of the adhesion *A* (area per node of the FE mesh), at that node,

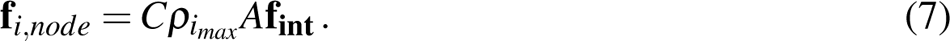

The fraction of *α*_5_*β*_1_-FN bonds *C* needs to be updated in time. For a given node,*i* given the previous value of the bond concentration, *C*, the updated bond concentration *C_t_*_+Δ*t*_ at each subsequent time step is based on the update

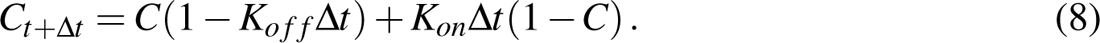

Note that the update eq. (8) is based on treating the bond kinetics in the limit of an ordinary differential equation discretized in time with an explicit Euler scheme.

The internal force balance for the cell and substrate include elastic deformation of the cell (*σ***_c_ ^pas^**), active contractile stress within the cell (*σ***_c_ ^act^**), and elastic deformation of the substrate (*σ***_c_ ^pas^**):

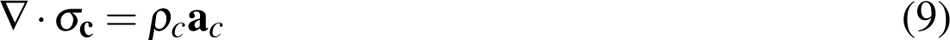

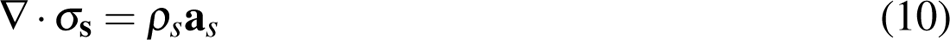

where *σ***_c_** = *σ***_c_^pas^** + *σ***_s_^act^** is the total stress in the cell, *σ***_s_** = *σ* **^pas^** is the total stress in the substrate, *ρ_c_*, *ρ_s_*are the densities of cell and substrate respectively (Table I), and **a***_c_,* **a***_s_*the corresponding accelerations.

The strong forms of the elastodynamic equations 9 and 10 have boundary conditions of the form *σ ·* **n** = **t** on boundary Γ. The strong forms are not directly evaluated. Rather, the internal forces were computed through the weak form. We multiplied both elastodynamic equations separately by test function, *ν*, integrated over a domain Ω of thickness 1*µ*m, and applied divergence theorem to get the following weak form for the cell (subscript c) and substrate (subscript s), respectively.

The *δ* **d** is the variation of the symmetric velocity gradient, i.e. virtual work by moving each node by an independent variation *δν*. **R** is the residual and the external force acting at a particular node of the respective cell and substrate meshes is:

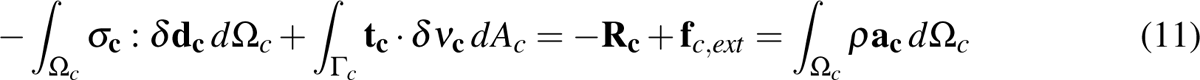

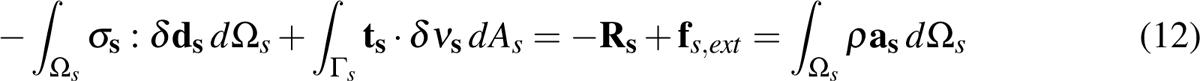

where **f***_i,node_* is the force due to integrin at each node, **f***_d_* is viscous drag, **f***_κ_* is curvature regularization, **f***_ac_* is a random fluctuation at the cell boundary from actin polymerization, and **f***_A_* is an area penalty to counteract cell contractility. Note that the nodal integrin force acts on the cell and substrate surfaces in opposite directions. The remaining variables act on the cell border and are further explained in the Supplementary Materials.

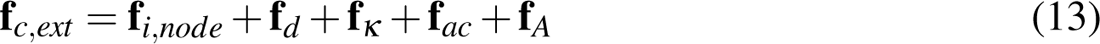

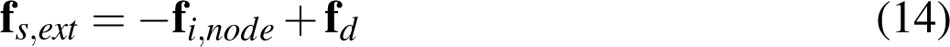

A dynamic explicit mesh generator, El Topo^29^, created and maintained the mesh during the simulation run. The explicit mid-point rule was used for time integration of the second order system of equations to update nodal velocities and positions. Three *α*_5_*β*_1_-FN stiffness values (*k_int_*) were used: 1pN/nm, 31pN/nm, and variable stiffnesses extracted from the MD simulation force-extension curves (MD-driven). The variable stiffness of the *α*_5_*β*_1_-FN complex within the FE model was modeled as a nonlinear spring by applying piece-wise linear interpolation in Python to the force-extension curves provided by the MD simulations as described in Section II D. Settings for each simulation run can be found in Supplementary Materials.

The overall sequence of the multiscale model is summarized in Figure 4. To summarize, the whole-cell FE model first imports the cell and substrate meshes and calculates the velocities and positions of the nodes. The *α*_5_*β*_1_-FN bonds are spread out uniformly across the surface of the cell with bond attachment points on the cell and the substrate. The displacement between the cell and substrate attachment points dictate the bond stretch. For the MD-driven case, the bond stiffness, *k_int_* is assigned based on the bond stretch. Otherwise, the stiffness is directly assigned according to each constant case (*k_int_*=1 or 31 pN/nm). The force per bond is then calculated via Hooke’s Law (eq. 6). This force is then used to update two things: the force per node (eq. 7) and the bond kinetics (eqs. 5 and 8). Cell contraction (eqs. 3 and 4) is then applied and the residual is computed via the weak form (eq. 11 and 12) considering the cell and substrate respective material properties (eq. 1 and 2), their elastodynamics (eqs. 9 and 10), and their force balances (eqs. 13 and 14). The nodal strains, velocities, and positions are updated and lastly, the simulation frame is saved. The whole-cell FE simulation iterates with a 1000-element mesh and a timestep of *dt* = 50*µs* over the course of an assigned time, *t_sim_* = 12*s*. Mesh and timestep convergence data can be found in the Supplementary Materials.

### D. Multiscale Model Coupling

The Gromacs function, `mdrun` outputted the force on the *α*_5_*β*_1_-FN complex. Furthermore, `gmx trajectory` was used to extract the center-of-mass coordinates of the restraints, LYS559 and GLU36, as well as the pull residue, PRO1142. The *α*_5_*β*_1_-FN extension length was measured in Python as the average vertical distance between PRO1142 and each of the two restrained residues. The resulting force-extension curve for each simulation run was then plotted. The `optimize` function from the `SciPy` library was used to produce a 5-segment piecewise linear fit on the 1 and 10 nm/ns force-extension curves, respectively. Ultimately, the 1 nm/ns curve-fit was used as a variable displacement-dependent spring constant in the whole-cell model to make up the “MD-driven” *α*_5_*β*_1_-FN stiffness, *k_int_*.

## III. RESULTS AND DISCUSSION

### A. ***α*_5_*β*_1_**-FN exhibited nonlinear and rate dependent stretching behavior under applied constant velocity

Prior to running the steered MD simulations at two pulling rates, the model’s energy minimized to −1.37e7 kJ/mol and the RMSD of the system plateaued while the pressure and temperature also remained stable during the NPT simulation (Supplementary Material). We chose 1 and 10 nm/ns pull rates for the steered MD simulations based on similar rates in other integrin subtypes^30,31^. As expected, *α*_5_*β*_1_-FN exhibited rate-dependent stretching behavior, meaning that the *α*_5_*β*_1_-FN force-displacement curves varied by pull rate (Fig. 2 A). The 10 nm/ns simulation reached a higher peak force of 723 pN and greater initial slope of 56 pN/nm compared to 444 pN and 31 pN/nm, respectively for the 1 nm/ns simulation.

**FIG. 2.**
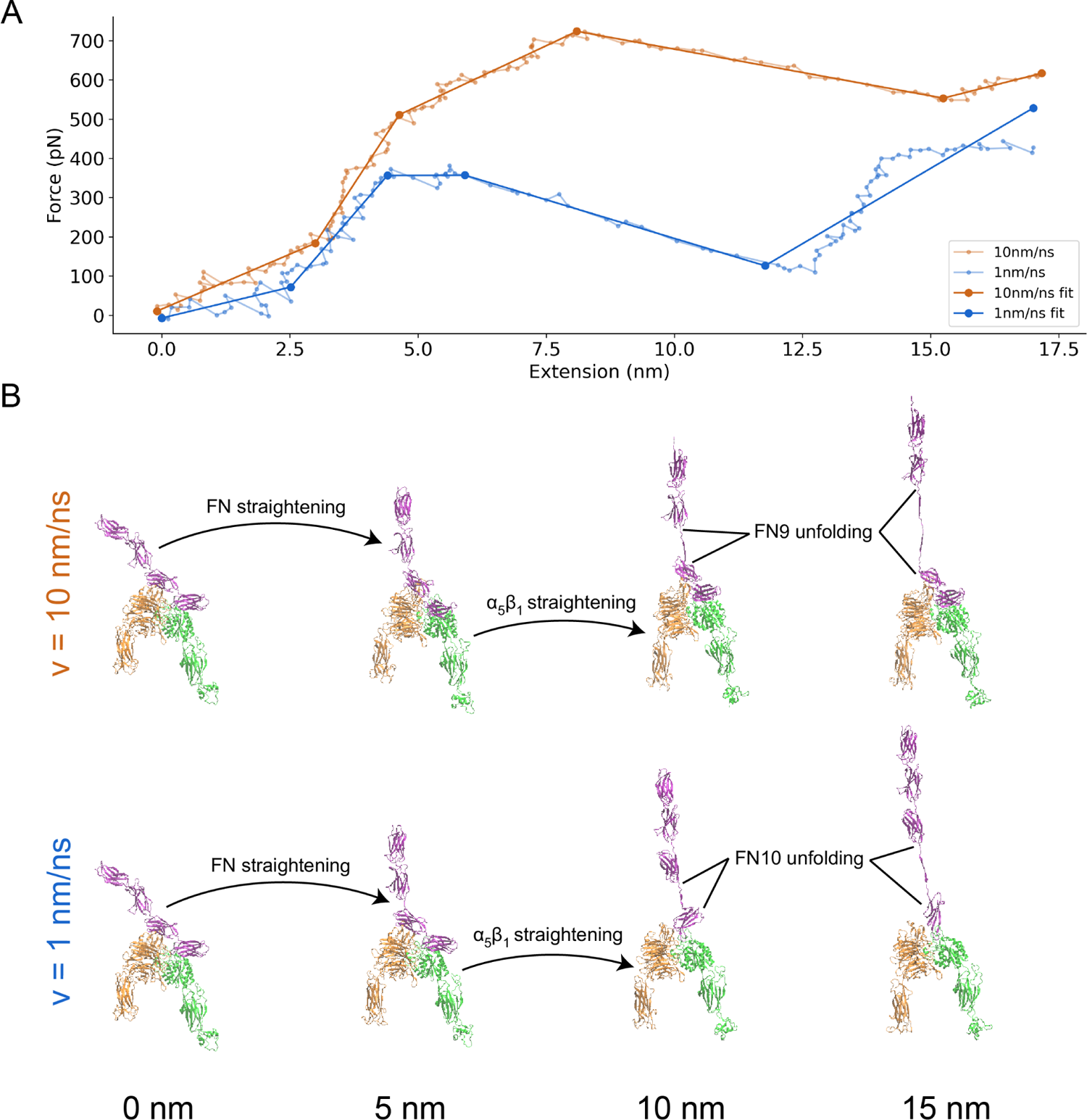
(A) Force-extension curve of *α*_5_*β*_1_-FN stretching at 10 and 1 nm/ns. The raw data are shown in transparent solid lines and the 5-segment piecewise linear fits are shown in opaque solid lines. (B) Frames of *α*_5_*β*_1_-FN during extension at 10 nm/ns and 1 nm/ns showing distinct stretching configurations at 0, 5, 10, and 15 nm of extension. In both cases, FN and *α*_5_*β*_1_ straightened before FN unfolded. However, for the 10 nm/ns case, the FN9 subdomain unfolded. Whereas for the 1 nm/ns case, FN10 unfolded. Movies showing *α*_5_*β*_1_-FN extension can be found in the Supplementary Materials.

In both cases, the stretching was dominated by FN, while integrin remained mostly rigid with some minor rotation and straightening. Curiously, at the faster 10 nm/ns pull rate, FN9 unraveled first before unbinding from the *α*_5_ head at the synergy site, whereas limited unraveling of FN was observed prior to unbinding for the slower 1 nm/ns pull rate (Fig. 2 B). Following the disconnection between FN and *α*_5_ at the synergy site, the force on the whole *α*_5_*β*_1_ integrin head became biased towards the RGD motif, causing the integrin heads to straighten with elongation of *α*_5_ and *β*_1_. However, the degree of head straightening was not consistent for both pull rates over the course of *α*_5_*β*_1_-FN extension. We opted to use radius of gyration (*R_g_*) as a proxy for integrin head straightness, with a larger radius indicating a straighter head. Visually, each integrin head started in a more closed positions with a relatively small *R_g_* before opening. Therefore, we believed it was appropriate to assume that a larger *R_g_* corresponded to a straighter molecule. For both rates, we observed an initial increase in the *R_g_* of both integrin heads prior to the unbinding of the salt bridge between arginine (ARG) 1379 in FN9 and aspartic acid (ASP) 154 in *α*_5_ (Fig. 5). However, the faster rate showed a sharp increase in *R_g_*of both heads after the salt bridge break at 6.1 nm, indicative of additional bonds pinning FN9 to *α*_5_ that then led to FN9 unfolding and *α*_5_ and *β*_1_ head straightening. In contrast, at the slower rate, we noticed a steady decrease in *R_g_* of both heads as FN10 unfolded immediately after the ARG1379-ASP154 break at 5.7 nm, presumably because *α*_5_ was allowed to relax after the departure of FN9. The faster rate elicits a greater reaction force out of *α*_5_*β*_1_-FN, which were resisted by other bonds between FN9 and *α*_5_ and a straightening of the integrin heads. This result was notable because it provided insight into how integrin may exhibit increased bond lifetime at higher forces, characteristic of previously observed catch bond behavior of integrins^15,32^.

**FIG. 3.**
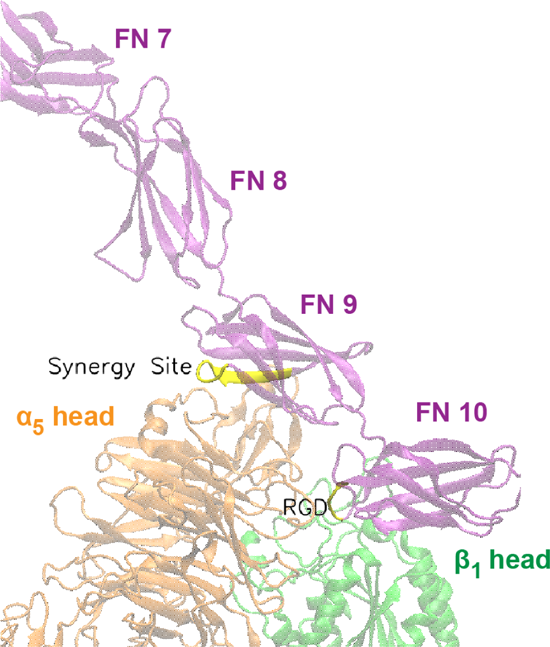
Close up view of DRVPHSRN synergy site and RGD motif/loop (shown in yellow) in FN that interact with the *α*_5_ and *β*_1_ heads, respectively.

**FIG. 4.**
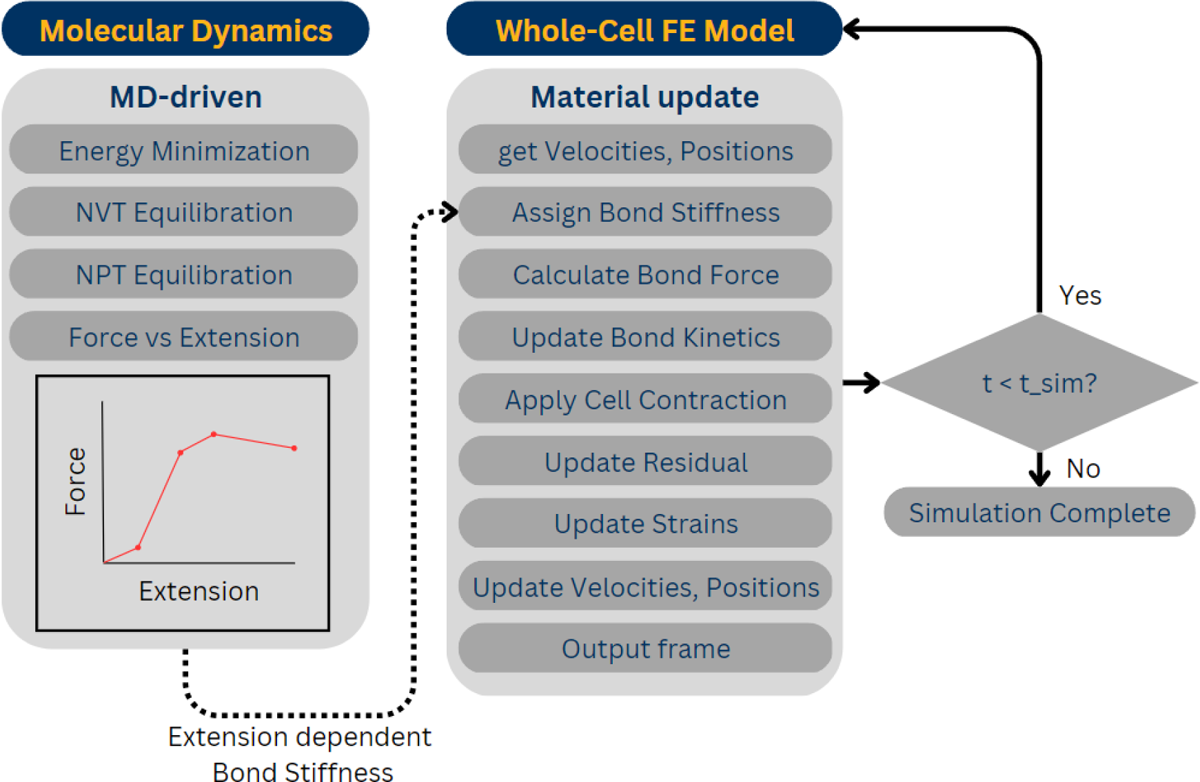
Multiscale framework that links the MD model to the FE model via a variable spring constant.

**FIG. 5.**
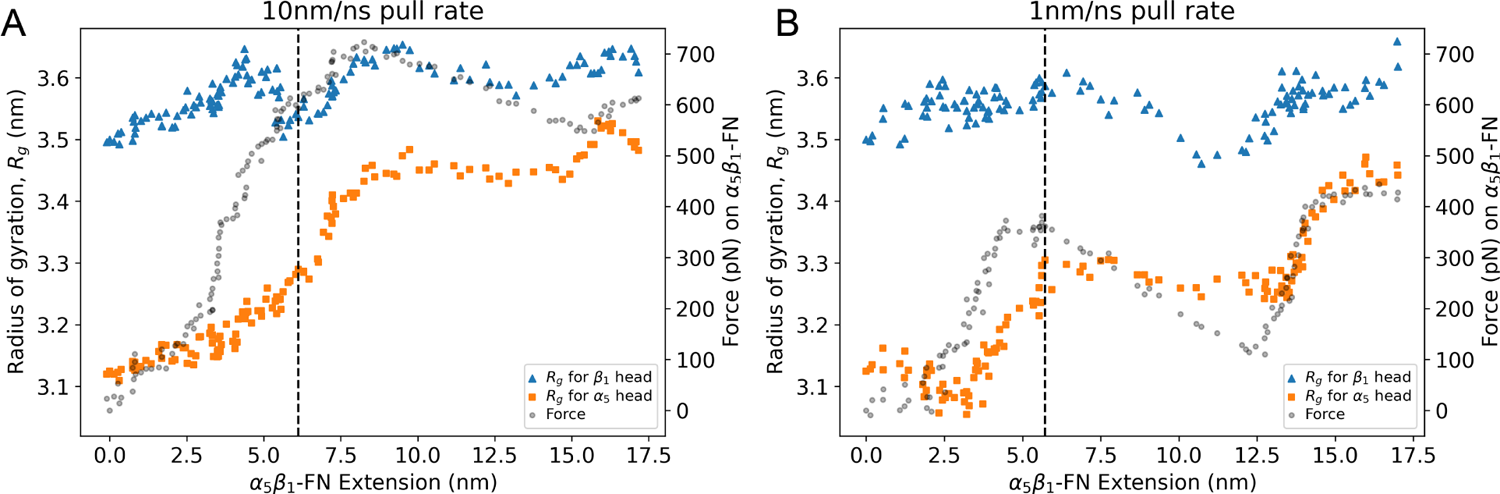
Radius of gyration (left vertical axis) of *α*_5_ and *β*_1_ heads and force (right vertical axis) on *α*_5_*β*_1_-FN during A) 10 nm/ns and B) 1 nm/ns extension. The dashed vertical line on each plot represents the moment the ARG1379-ASP154 salt bridge was broken.

The observed viscoelastic behavior of *α*_5_*β*_1_ has been shown both experimentally and computationally. Single-molecule AFM studies show higher rupture forces at faster pull rates^20^ and separate steered MD simulations of integrin^30,31^ and FN^33^ showed rate-dependent and force-dependent stretching behavior seen in viscoelastic materials. We expected this viscoelastic behavior to remain when *α*_5_*β*_1_ and FN are in complex. To confirm, we tested *α*_5_*β*_1_-FN’s viscoelasticity *in silico* via constant force simulations at 300 and 500 pN, similar to what would be done in a mechanical creep test where constant stress is applied (Fig. 6). We fit the Bausch viscoelasticity model, which combines a Kelvin model with a dashpot in series^34^, to the extension-time plots, supporting the characterization of *α*_5_*β*_1_-FN’s time-dependent stretching and viscoelastic nature.

**FIG. 6.**
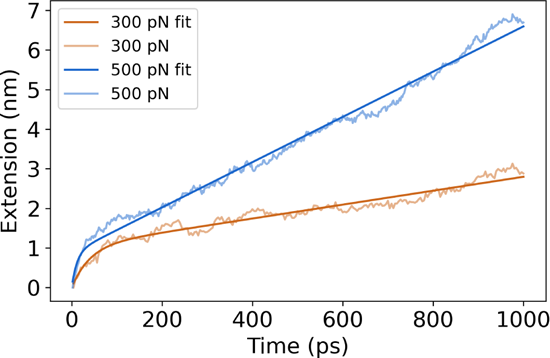
Extension plots of constant force simulations at 300pN and 500pN pulling forces. The Bausch^34^ viscoelastic model was fit to each of the plots.

While our MD simulations and previous literature have demonstrated the nonlinear stretching behavior of *α*_5_*β*_1_-FN, multiscale models assume a linear integrin stiffness between 0.001-2 pN/nm^27,28,35^. Recent multiscale models have used this assumption when analyzing fundamental phenomena such as integrin activation, organization, and clustering at the cell and tissue scales^27,28,35^. Most recently, Guo et al. showed a framework that combined adhesion kinetics with the finite element method (FEM) to model stretch-driven mechanosensing at the tissue level by coupling integrin adhesion with the nonlinear tissue mechanics of fibrin and collagen^27^. While these models provide unique insights into multiscale mechanobiology of cell adhesion, for models to account for integrin and FN’s nonlinear stretching behavior, a dynamic spring stiffness that adjusts depending on extension is required. For our work, we used our steered MD force-extension plots to inform a dynamically changing spring in a continuum model of the whole cell.

A limitation of our approach is that MD simulations are computationally expensive and runtimes would be unreasonably long if we adopted experimentally relevant 800 nm/s pull rates used by past AFM studies^36,37^. However, using faster pull rates leads to higher single-molecule forces beyond 300pN as was noticed in our force-extension curves. Moreover, the MD model limited the flexibility of the proximal ends of the integrin heads by restraining them with a harmonic spring, potentially contributing to larger measured forces. The heads may have otherwise been more free to move depending on the motion of the integrin legs and tails within the cell membrane, which were not modeled to reduce computational cost and add model stability. Previous studies found average *in situ* rupture forces for *α*_5_*β*_1_-FN to be 34^36^ and 38.6 pN^37^ in endothelial cells and cardiomyocytes, respectively. Single molecule AFM conducted by Li et al. measured a mean rupture force of *α*_5_*β*_1_-FN of 69 pN at a loading rate of 1800-2000 pN/s, with a peak rupture force of 120 pN at 18,000 pN/s^20^. More recently, FRET-based sensors were used to measure adhesion forces between 1-7 pN on fibroblasts plated on glass^21^. All these measured forces are much lower than those predicted by the MD simulations. Higher forces at much faster pull rates meant that our *α*_5_*β*_1_-FN stiffness results were significantly larger than what has been observed *in vitro*. However, in all the experiments, the nonlinearity of *α*_5_*β*_1_-FN’s stretching behavior was apparent, challenging the linear stiffness assumption made by previous models^27,28,35^. Furthermore, while an average FN stiffness of 0.5 pN/nm has been reported^38,39^, the coupled *α*_5_*β*_1_-FN stiffness has not. Additionally, our steered MD simulations provided atomic level details that helped explain how key binding sites contributed to pull rate dependent nonlinear stretching.

### B. Force Distribution Analysis of ***α*_5_*β*_1_**-FN reveals dynamics of adhesion-mediating residues that contribute to nonlinear force-extension behavior

Visualization of the coulombic interactions via Force Distribution Analysis of the steered MD results demonstrated how key adhesion mediators could contribute to nonlinear, rate-dependent, force-extension of *α*_5_*β*_1_-FN. Two key mediators are the DRVPHSRN synergy site and the RGD motif in FN (Fig. 3). In our system, the FN synergy site was represented by residues 1373 to 1380 and the RGD motif was represented by residues 1493 to 1495. Spinning disk microscopy has previously shown that mutating one to two select residues on the synergy site leads to a decrease in overall cell adhesion and mutating the RGD motif eliminates cell adhesion force completely^15^. Furthermore, inducing a synergy site mutation or an RGD deletion leads to a reduction in single molecule rupture force of *α*_5_*β*_1_-FN^20^. Therefore, we looked closely at the dynamics of these adhesion mediators during *α*_5_*β*_1_-FN stretching at 1 nm/ns and 10 nm/ns.

Interestingly, the *α*_5_*β*_1_-FN extension showed two modes of stretching depending on the pull rate. Heatmaps overlaid on the molecule illustrated the degree of coulombic interaction, where “hotter” or “redder” zones indicated larger pairwise punctual stresses. For the 10 nm/ns case, the ARG1379-ASP154 salt bridge is broken after 6.1 nm of *α*_5_*β*_1_-FN extension (Fig. 7A). This action then loosens the grip between *α*_5_ and FN9, allowing FN9 to rotate to find a new interaction between glutamic acid (GLU) 1405 and serine (SER) 85. FN9 then unfolded, contributing to the initial decrease in force and most of the extension before GLU1405 and SER85 release. Between 0 and 5 nm, FN began to straighten while simultaneously tugging on the synergy site and RGD. The force-extension response “softened” as the salt bridge was broken and FN9 started to rotate. The large extension and reduction in force past 8 nm (Fig. 2) was due to the rapid unfolding of FN9 while GLU1405-SER85 pinned the rest of FN9 in place. After two strands of FN9 are unwound, the applied load became directed at the GLU1405-SER85 pin until it finally separated. Notably, the unfolding pathway with two strands unwound of FN9 has been illustrated before in constant force simulations of FN^33^. Our model corroborates these results while providing new insight into the dynamics of FN unfolding when interacting with *α*_5_*β*_1_ integrin.

**FIG. 7.**
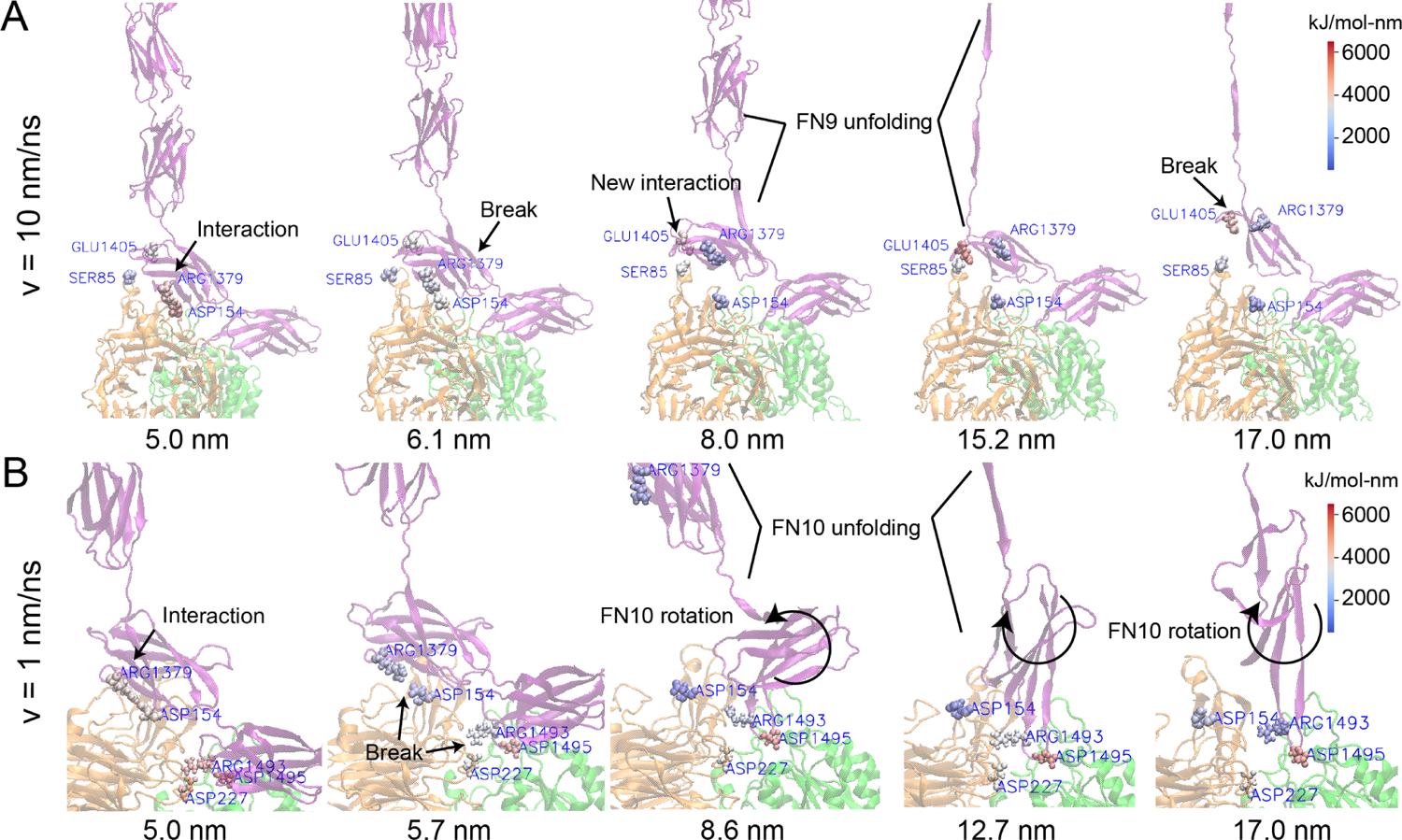
Force Distribution Analysis of *α*_5_*β*_1_-FN for two pull rates at key events. The color map refers to the punctual stress (in kJ/mol-nm) at each residue. (A) At 10 nm/ns, there was a coulombic interaction at the ARG1379-ASP154 salt bridge and no interaction between GLU1405 and SER85. As FN was extended, the salt bridge ruptured and allowed FN to rotate and establish a new interaction between GLU1405 and SER85. FN9 continued to unfold, increasing stress on the GLU1405-SER85 connection, eventually breaking it. (B) At 1 nm/ns, the ARG1379-ASP154 salt bridge, part of the synergy site, together with ARG1493 and ARG1495, part of the RGD motif, maintained a hold on FN. As FN extended, increased stress led to the simultaneous rupture of ARG1493-ASP227 and ARG1379-ASP154. This allowed FN10 to unfold and rotate. ARG1493-ASP227 disconnected and reconnected throughout the remainder of the simulation. Movies showing the FDA can be found in the Supplementary Materials.

The observed unbinding and unfolding sequence in *α*_5_*β*_1_-FN was not preserved at 1 nm/ns. The salt bridges, ARG1379-ASP154 and ARG1493-ASP227 simultaneously broke at 5.7 nm of extension after a short force plateau between 4.8-5.7nm, but unlike in the 10nm/ns run, FN9 did not create a new interaction with *α*_5_ (Fig. 7B). Rather, FN10 unfolded, leading to the majority of the overall extension and reduction in force from 5.7-12.7nm (Fig. 2A). During FN10 unfolding, the interaction between ARG1493 in FN and ASP227 in *α*_5_ alternated between high and low coulombic interactions while ARG1495 maintained adhesion with *β*_1_ integrin. Due to the lack of interaction between the synergy site in FN9 and *α*_5_, FN9 was free to separate from integrin so FN10 could readily unfold. Once one strand had completely unfolded, due to the direction of the pulling force with respect to the orientation of FN10, the force needed to rotate FN10 prior to unwinding the second strand, which led to an increase in force (Fig. 2B).

At both pull rates, the synergy site and RGD loop played key roles in maintaining the adhesion between *α*_5_*β*_1_ and FN. Specifically, the salt bridge between ARG1379 and ASP154 contributed to the molecule’s initial “stiff” behavior prior to FN unfolding; and part of the RGD loop between *β*_1_ and FN10 was the only remaining connection between integrin and FN after full extension. Due to their instrumental role, it stands to reason that interfering with these residues via point mutations would reduce adhesion^15^ and rupture force^20^. While measured *in vitro* forces on *α*_5_*β*_1_-FN have been shown to be much smaller than we have presented due to our model’s much faster pulling speed, nonlinear force-extension behavior and rapid jumps in force have been observed^15,20,21^. We showed how key residues could contribute to this characteristic behavior during *α*_5_*β*_1_-FN extension in a pull rate dependent manner. To bridge the nanoscale integrin stretching to cell-scale integrin dynamics, as a proof-of-concept, we modeled the force-extension of *α*_5_*β*_1_ as a nonlinear spring and used it to scale up to a 2D whole-cell continuum model.

### C. Multiscale integration of ***α*_5_*β*_1_**-FN force-extension with whole-cell integrin dynamics

Prior to integrating the force-extension curves from the MD runs, we had ran a baseline simulation of the whole-cell model with similar parameters to those commonly used in literature^27,28,35^. In particular, we set the *α*_5_*β*_1_-FN stiffness, *k_int_*, to 1pN/nm. For all simulations, the cell contractility was ramped from 0 to 200Pa within the first 2s and held at 200Pa for the remainder of the 12s simulation. Integrins were recruited to the cell border, achieving maximum concentration and force as the contractility reached 200Pa at 2s (Fig. 8).

**FIG. 8.**
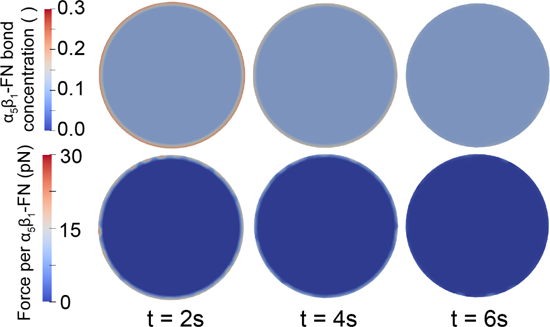
The dimensionless *α*_5_*β*_1_-FN bond concentration (top) and force per *α*_5_*β*_1_-FN (bottom) results for the baseline whole-cell simulation with *k_int_*= 1pN/nm. *α*_5_*β*_1_-FN localization and force dissipation occurred rapidly and no significant changes in distribution were observed past 6s. Movies showing simulation trajectories can be found in the Supplementary Materials.

Integrin’s spatial distribution on the cell’s leading edge during motion has been previously observed *in vitro*^22^, corroborating the results from the model. During contraction, the model’s average peak bond concentration reached 10.6% (Fig. 9A) with a max peak of 22.5%. The average force per bond followed a similar curve, reaching an average peak of 1.9pN (Fig. 9B) with a max peak of 28.6pN at the cell boundary. These bonds had short lifetimes and dissociated quickly, allowing the model to dissipate the contraction and reach equilibrium just before the 6s mark. After reaching this equilibrium point, the mean force was 0.17*±*0.04pN with max forces reaching 15.9pN at the boundary. The peak bond forces and concentrations occurred on the boundary due to the positive feedback loop of the catch-slip bond dynamics. While the strain across the cell is uniform due to the applied isotropic contractility, the deformation of the bond springs are the greatest at the boundary, leading to higher bond concentrations and forces. Overall, the forces were within the 1-38pN range that has been observed *in vitro*^21,36,37^ and well within the peak single *α*_5_*β*_1_-FN rupture forces measured via AFM of 120pN^20^.

**FIG. 9.**
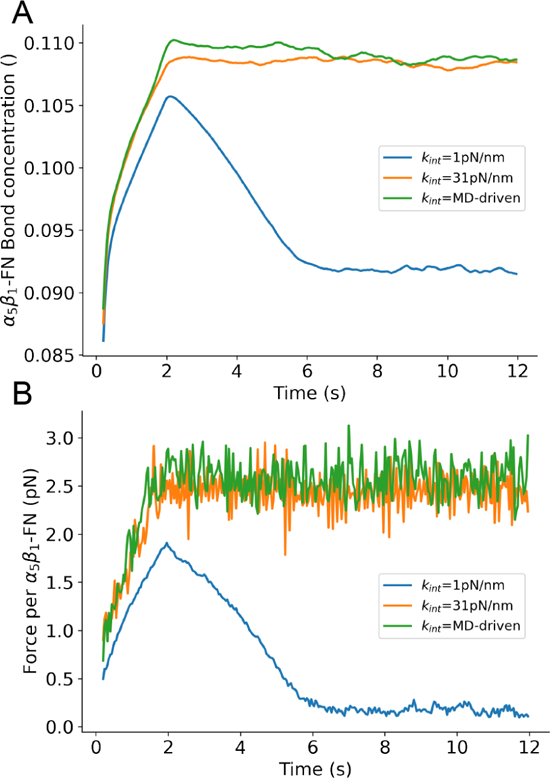
Whole-cell average A) *α*_5_*β*_1_-FN bond concentration (dimensionless) and B) force per *α*_5_*β*_1_-FN over the simulation run. Three test conditions for *α*_5_*β*_1_-FN stiffness are shown per plot: 1) constant 1pN/nm baseline from past models^27,28,35^, 2) constant 31pN/nm based on the first segment of the 1nm/ns force-extension curve fit, and 3) MD-driven stiffness derived from using all segments of the curve fit.

The baseline simulation provided a control to test against our two simulation conditions derived from the 1 nm/ns MD simulation. We defined a varying, MD-driven *α*_5_*β*_1_-FN stiffness as the entire 1nm/ns force-extension curve fit. To evaluate how the nonlinearity of the MD-driven integrin spring affected whole-cell adhesion dynamics, we used the slope of the first segment, 31pN/nm, to define a constant *α*_5_*β*_1_-FN stiffness test condition.

Overall, the *α*_5_*β*_1_-FN bond concentration for the constant and MD-driven *α*_5_*β*_1_-FN stiffness conditions followed a similar trend and were both slower to distribute the contraction load (Fig. 9) than the 1pN stiffness setting. Past 2s, the mean forces steadied at 2.45*±*0.18pN and 2.59*±*0.19pN for the constant and MD-driven runs, respectively. The noise in the the bond concentrations and force per bond (Fig. 9) were due to the random 5pN actin polymerization force. The results for both cases were similar. The constant 31pN/nm run reached a max average bond concentration of 10.9% and the MD-driven case topped at 11.0%. Max average forces, located at the cell boundary (Fig. 10), were 53.5pN and 55.6pN for the 31pN/nm and MD-driven runs, respectively. The positive feedback loop of the catch-slip bond at the boundary continued to drive the peak forces and concentrations across all stiffness settings.

**FIG. 10.**
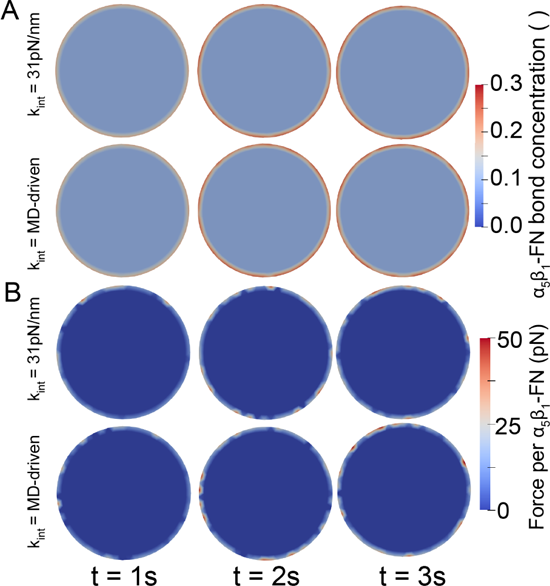
Whole-cell simulation results for the constant and MD-driven spring stiffnesses. A) *α*_5_*β*_1_-FN bond concentration and B) Force per *α*_5_*β*_1_ integrin at three time frames within the first 3s of the simulation. Dissipation continued past 3s, but the changes were minor. Movies showing simulations can be found in the Supplementary Materials.

Notably, model predictions surpass *in situ* rupture forces of 34-38.6pN for *α*_5_*β*_1_-FN^36,37^ and 40pN for another subtype, *α_V_ β*_3_^40^. Chang et al. used FRET-based sensors to measure adhesion forces between 1-7 pN on fibroblasts^21^. Recent work has used leveraged tension gauge tethers to measure single molecule forces on RGD-binding integrins and showed that integrin activation occurs below 12 pN and *α_V_ β*_1_ could sustain forces over 54pN in mature FAs^41^. In summary, the models we presented showed estimations towards the upper bounds of measured biophysical forces felt by integrin.

The MD-driven and constant 31pN/nm integrin stiffness models showed similar force and concentration results indicating that linear spring stiffness was sufficient to capture *α*_5_*β*_1_-FN molecular dynamics in this model. Notably, bond lengths were maintained below 2.5nm, where the stiffness jumps to 99.5pN/nm in the MD-driven force-extension curve (Fig. 2A). The main difference observed in the bond force and concentration response was between soft (1pN/nm) and stiff (31pN/nm, MD-driven) integrin models. These differences arose due to the force balance between the cell, the substrate, the integrin, and other random forces (eq. 13). All these forces contributed to the integrin deformation, *u_int_* (Fig. 11), which was multiplied by integrin stiffness to calculate force. This bond concentration was updated based on this bond force and catch-slip bond model (Fig. 11 and eq. 8). In our case, the bond lengths ranged from 0-15.9nm for the soft integrin and 0-1.8nm in the stiff integrin. This led to forces between 0-15.9pN and 0-55.6pN for the soft and stiff integrin, respectively. To summarize, the balance between applied forces, cell/substrate material properties, and integrin stiffness led to varying bond deformation which contributed to alterations in bond concentration due to catch-slip bond dynamics.

**FIG. 11.**
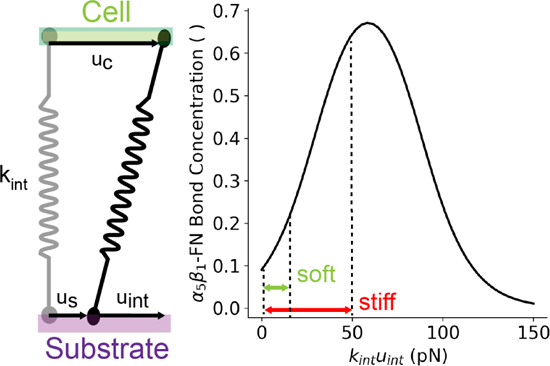
Schematic of the balance at an equilibrium state between the cell, substrate, and spring deformations contribute to changing bond concentrations based on the catch-slip bond curve (eq. 8).

## IV. CONCLUSION

We developed a coupled multiscale model which showed how amino acid interactions at the synergy site in FN contribute to the nonlinear force-extension behavior of *α*_5_*β*_1_-FN, which leads to unique whole-cell adhesion force landscapes. The model demonstrated whole-cell integrin spatial distribution along the cell membrane, consistent with fibroblasts plated *in vitro*^22^ and forces within the 120pN maximum single molecule rupture force and 1-38 pN *in situ* rupture forces^21,36,37^.

This study has limitations. We used high pull rates in the MD simulations to maintain reasonable computational runtimes. However, this led to large forces during *α*_5_*β*_1_-FN extension. While the computational cost is a common drawback of MD, the detailed data and outputs gained from the amino acid dynamics and their connection to whole-cell integrin dynamics would have been otherwise unobservable. Therefore, we believe that it was useful to include this demanding piece of the multiscale model. A combination of slower pull rates and coarse grained MD simulations could be the compromise necessary to investigate the nonlinear mechanics while maintaining some nanoscale details.

Also, we chose *α*_5_*β*_1_ integrin as the sole surface receptor, but cells have additional subtypes with varying roles^35,42^ and potentially different adhesion strengths^41^ and binding kinetics^43,44^. Given the 24 known subtypes of integrin^45^, it is critical to understand which ones are the main contributors to adhesion maintenance in the presence of specific ligands. For example, in the case of fibronectin, a recent single-cell force spectroscopy study indicated that pan integrin knockout fibroblasts only expressing *α*_5_*β*_1_ and *α_V_ β*_3_ transmitted the same amount adhesive force as wildtype fibroblasts on a fibronectin coated surface^46^. Therefore, extending our model to contain these two subtypes may be an appropriate approximation to evaluate integrin adhesion mechanics for fibroblasts on fibronectin. Another key consideration is the dynamics of low-affinity and high-affinity conformations of integrin. For our model, we assumed that *α*_5_*β*_1_ integrin was in a high affinity, extended-open conformation. However, it has been demonstrated that low-affinity bent-closed and extended-closed conformations of *α*_5_*β*_1_ and *α_V_ β*_3_ can still bind to fibronectin^44,47^. To include the contributions of varying subtypes, it would be necessary to update to our catch-slip bond model (Fig. 11) to include high and low affinity conformational states, manage the population distribution of integrin subtypes as done in other models^28,35^, and expand on existing steered MD characterizations of *α_V_ β*_3_^30,48^ to add to ours of *α*_5_*β*_1_. Overall, more investigation is needed to evaluate how integrin subtypes collaborate to manage cell adhesion dynamics.

The model assumed a homogeneous substrate. However, tissue microenvironments are spatially heterogeneous and respond to the binding and unbinding dynamics between ECM fibers^49–53^. This leads to viscoplastic material behavior, or time and frequency dependent force dissipation^54^ which mediates cell migration, differentiation, and disease progression^55–57^. To include these effects, we could represent the substrate viscoplasticity via the Norton-Hoff constitutive model^49,58^, and the cell’s myosin-actin engagement via the molecular or motor clutch model^53,59^. We would expect a heterogeneity to arise in the force and spatial distribution of the integrin bonds, localizing near denser packs of crosslinked fibers. We hypothesize that stiffer integrins would lead to denser packing of ECM fibers due to their slow rate of sustained force compared to softer bonds. However, more investigation is needed to reveal the relationship between cell adhesion and force-mediated ECM fiber mechanics.

Our model focused on cell adhesion mechanics and has the potential to grow into a framework that can investigate cell mechanotransduction across multiple scales. For example, we could test how unique mutations on integrins affect whole-cell dynamics *in silico*. Additionally, by incorporating the cell nucleus, we could support early evidence to show how its mechanosensitive nature and material properties could govern gene transcription^60–62^. Key components that have previously been modeled such as the cell membrane, integrin’s transmembrane domain, and integrin clustering and diffusion^28,35,63–65^ were omitted from our model for simplicity, but could be added as new multiscale mechanobiological questions are posed regarding their mechanics.

Lastly, our multiscale framework could be broadened to reveal the nano- and micro-mechanics within nascent engineered tissues and organ-chips that apply controllable biophysical loads at the cell membrane^66–71^.

## Supporting information

Movies of Simulation Trajectories

## SUPPLEMENTARY MATERIAL

See the supplementary material for detailed equations and parameters for the whole-cell model; parameters for the minimization, equilibration, and force distribution analysis of the MD model; RMSD, pressure, and temperature during equilibration; whole-cell model mesh and timestep convergence studies; and trajectory movies for the whole-cell and MD models. The whole-cell model is available at https://github.com/dredremontes/wholeCellFE and the MD model is available at https://github.com/dredremontes/pull_integrinMD.

## AUTHOR CONTRIBUTIONS

**A.R. Montes**: Conceptualization, data curation, formal analysis, funding acquisition, investigation, methodology, project administration, software, validation, visualization, and writing - original. **G. Gutierrez**: Formal analysis, investigation. **A.B. Tepole**: Conceptualization, data curation, formal analysis, funding acquisition, investigation, methodology, project administration, resources, software, supervision, validation, visualization, writing - original, and writing - review & editing. **M.R.K. Mofrad**: Conceptualization, funding acquisition, project administration, resources, supervision, and writing - review & editing.

## ACKNOWLEDGMENTS

This research used Stampede2 at Texas Advanced Computing Center through allocation MCB100146 from Advanced Cyberinfrastructure Coordination Ecosystem: Services & Support (ACCESS) super-computing facilities, supported by the National Science Foundation grants #2138259, #2138286, #2138307, #2137603, and #2138296. A.R.M. was funded by the Ford Foundation’s Predoctoral Fellowship awarded by the National Academies of Science, Engineering, and Medicine and the Robert N. Noyce Fellowship from UC Berkeley’s College of Engineering. G.G. was supported by the National Science Foundation California Alliance for Minority Participation. Thank you to Ghafar Yerima and Nya Domkam for helping set up GROMACS and gromacs-fda. We also thank the Molecular Cell Biomechanics Lab and the Berkeley Biomechanics Lab for fruitful discussions that improved the manuscript.

## DATA AVAILABILITY STATEMENT

The data that support the findings of this study are available from the corresponding author upon reasonable request.

