## Supplementary Material for "Multiscale Computational Framework to Investigate Integrin Mechanosensing and Cell Adhesion"

**Author's note**: The parameters used for the simulations are listed in the following tables for reference. The code for the molecular dynamics (MD) simulations and whole-cell finite element (FE) simulations are available on Github.

- `https://github.com/dredremontes/pull_integrinMD`
- `https://github.com/dredremontes/wholeCellFE`

The supplementary material contains:

- Finite Element (Whole-cell) Model Equations
- Table S1: Energy Minimization Parameters
- Table S2: NVT Parameters
- Table S3: NPT Parameters
- Table S4: Steered MD Parameters
- Table S5: Force Distribution Analysis Parameters
- Table S6: Whole-cell model parameters
- Fig. S1: Energy Minimization and NVT RMSD
- Fig. S2: NPT RMSD, Pressure, and Temperature
- Fig. S3: Whole-cell Model Meshes
- Fig. S4: Whole-cell Model Mesh Convergence
- Fig. S5: Whole-cell Model Timestep Convergence
- Movie S1: 1nm/ns extension of $\alpha_5\beta_1$-FN
- Movie S2: 10nm/ns extension of $\alpha_5\beta_1$-FN
- Movie S3: 1nm/ns Force Distribution Analysis
- Movie S4: 10nm/ns Force Distribution Analysis
- Movie S5: All Whole-cell simulations

The movies are in separate .mp4 files.

**Finite Element (Whole-cell) Model Equations**

The triangular mesh is handled through the object `SurfTrack` part of the ElTopo library `https://github.com/tysonbrochu/eltopo`. Two `SurfTrack` objects are created, one for the cell mesh and one for the substrate mesh. The `SurfTrack` contains the nodal coordinates which we denote $\mathbf{x}_t$, the connectivity `tri_mesh`, and an additional connectivity which for every node stores the *one ring*, i.e. the nodes adjacent to a given node, which we refer to as `node_onering`. There is also a flag for nodes on the boundary of the surface. We modified the original library to also store nodal velocities, accelerations, and previous value of the strain. These vectors are named $\mathbf{v_t}$, $\mathbf{a}_t$, $\mathbf{e}_t$. For the cell mesh we also define one more vector field associated with the nodes, $\mathbf{u}_i$, that contains the displacement vector of an integrin-ligand bond with respect to its stress-free state. Lastly, for the cell mesh we also store a scalar field $C$ with the local integrin-bound fraction as described in the main text.

The integration is done explicitly with the midpoint rule. Given the current value of positions, velocities and accelerations, the midpoint velocity is calculated as

$$\mathbf{V}_{t+0.5\Delta t} = \mathbf{V}_t + \frac{\Delta t}{2}\mathbf{A}_t \,. \tag{S1}$$

Then the the updated positions are computed as

$$\mathbf{X}_{t+\Delta t} = \mathbf{X}_t + \Delta t \mathbf{V}_{t+0.5\Delta t} \,. \tag{S2}$$

Given the updated positions, the forces at the nodes are computed with the weak form. As described in the main text, the residual of internal forces is

$$\mathbf{R} = -\int_{\Omega} \sigma : \delta\mathbf{d}\, d\Omega \tag{S3}$$

Computation of eq. (S3) is done with standard linear triangular finite element interpolation. The residual is calculated independently for the cell and substrate meshes. Additionally, as noted in the main text, the cell has two components of the stress, a passive and an active component, whereas the substrate only has a passive component. The code is in an updated Lagrangian framework, whereby the strain at the previous time step $\mathbf{e}_t$ gets updated based on the updated nodal coordinates $\mathbf{x}_{t+\Delta t}$ by updating the left Cauchy-Green deformation

$$\mathbf{B}_{t+\Delta t} = \mathbf{F}\mathbf{b}_t\mathbf{F}^{\top} \,, \tag{S4}$$

where $\mathbf{F}$ denotes the deformation gradient from the incremental deformation $\mathbf{x}_t$ to $\mathbf{x}_{t+\Delta t}$.

The external force acting at a particular node of the cell mesh is

$$\mathbf{f}_{c,ext} = \mathbf{f}_{i,node} + \mathbf{f}_d + \mathbf{f}_\kappa + \mathbf{f}_{ac} + \mathbf{f}_A \,, \tag{S5}$$

where $\mathbf{f}_{i,node}$ is the force due to integrin at each node, $\mathbf{f}_d$ is viscous drag, $\mathbf{f}_\kappa$ is curvature regularization, $\mathbf{f}_{ac}$ is a random fluctuation at the cell boundary from actin polymerization, and $\mathbf{f}_A$ is an area penalty to counteract cell contractility. The force from integrin at each node is defined in the main text:

$$\mathbf{f}_{i,node} = C\rho_{i_{max}} A k_{int} \mathbf{u_{int}} \,. \tag{S6}$$

The viscous drag is

$$\mathbf{f}_d = -d\mathbf{u} \,, \tag{S7}$$

where $\mathbf{u} = \mathbf{x}_{t+\Delta t} - \mathbf{x}$ is the displacement of the node and $d = 0.001\,\text{pN}/\mu\text{m}$ is a small drag coefficient. The curvature force is calculated only for nodes on the boundary. Discrete curvature $\kappa$ is approximated based on twice the turning angle along the boundary curve divided by the length of the curve. Given the curvature, the force is

$$\mathbf{f}_\kappa = -k_\kappa \kappa \frac{l_b}{2}\mathbf{n}, \tag{S8}$$

where $\mathbf{n}$ is the outward unit normal at the boundary and $k_\kappa = 20\text{pN}/\mu\text{m}$ is a small bending stiffness to prevent buckling and ruffling of the boundary, and $l_b$ is the length of the edges incident at a node on the boundary. The actin polymerization fluctuation is also only at the boundary

$$\mathbf{f}_{ac} = f_{ac}\mathbf{n}, \tag{S9}$$

where $f_{ac}$ is a random force computed from sampling of a Poisson process. Specifically, an actin polymerization rate of $k_{ac} = 10\text{s}^{-1}$ was assumed [1]. The density of actin monomers near the boundary was estimated to be $\rho_{ac} \approx 100\text{monomer}/\mu\text{m}$ [2], and the individual monomer force to be 5pN [3]. To compute $f_{ac}$, we first determined the number of possible events at a node to be proportional to $n_{ac} = \rho_{ac} l_b/2$, where $l_b$ is the length of the edges incident to the boundary node. Then, we performed a `for` loop over $n_{ac}$ and for each iteration sampling a random number from $p_{ac}\ \mathscr{U}([0,1])$ and if $p_a c < k_{ac}\Delta t \exp(-k_{ac}\Delta t)$, we updated the actin force as $f_{ac} \leftarrow f_{ac} + 5\text{pN}$.

Finally, the area constraint is also only applied at the boundary nodes and it is

$$\mathbf{f}_A = -p_A(A_{tot} - A_0)\frac{l_b}{2}\mathbf{n}, \tag{S10}$$

where $A_{tot}$ is the entire area of the cell, $A_0$ is an attractor for the area, and the length of the curve associated with the node is $l_b$ as before, just as the normal is also associated with the node and it is the outward unit normal as before. The strength of this constraint is imposed with the pressure parameter $p_A = 1\text{pN}/\mu\text{m}$. In reality the effects of the regularization terms is small but allows to keep the simulation stable and correspond to physically meaningful phenomena.

For the substrate the only contributions are

$$\mathbf{f}_{s,ext} = -\mathbf{f}_{i,node} + \mathbf{f}_d\,. \tag{S11}$$

Note that the force from integrin acts in the opposite direction on the substrate compared to the cell. Given the force in the cell or the substrate, $\mathbf{f}_{ext}$, the acceleration is updated by:

$$\mathbf{a}_{t+\Delta t} = \mathbf{M}^{-1}(\mathbf{f}_{ext} - D\mathbf{M}\mathbf{v}_{t+0.5\Delta t}) \tag{S12}$$

with $\mathbf{M}$ being the respective diagonal mass matrix and a damping coefficient of $D = 0.001$ 1/s. Lastly, the velocities are updated

$$\mathbf{v}_{t+\Delta t} = \mathbf{v}_{t+0.5\Delta t} + \frac{\Delta t}{2}\mathbf{a}_{t+\Delta t}\,. \tag{S13}$$

One additional set of updates in the model is the change of the displacement of the integrin-ligand pair $\mathbf{u}_i$ as the cell and substrate deform. From the cell deformation, the update is

$$\mathbf{u}_i \leftarrow \mathbf{u}_i + \mathbf{u}_{cell}\,, \tag{S14}$$

where $\mathbf{u}_{cell} = \mathbf{x}_{t+\Delta t} - \mathbf{x}_t$ is nothing but the corresponding displacement of the node on the cell. However, for the substrate, the process is slightly more involved. Since the $\mathbf{u}_i$ is associated with the nodes of the cell mesh, we first get the triangle in the substrate mesh this corresponds to and then update

$$\mathbf{u}_i \leftarrow \mathbf{u}_i - \mathbf{u}_{subs}\,, \tag{S15}$$

where $\mathbf{u}_{subs}$ is the interpolated displacement of the substrate at the correct location of the node from the cell mesh. Because when forms break and new bonds form they are not assumed to be pre-strained, there is some dissipation associated with this drift in the reference configuration of the integrin-ligand bond stretch, captured by

$$\mathbf{u}_i \leftarrow \mathbf{u}_i(C_t - \Delta t K_{off} C_t)/C \tag{S16}$$

which reduces the stretch in the integrin proportional to the number of broken bonds with respect to the previous integrin-ligand bond fraction $C_t$. If $C_t - \Delta t K_{off} C_t \leq 0$ then the stretch of the integrin-ligand bonds is set to $\mathbf{0}$.

[3] Serge Dmitrieff and François Nédélec. Amplification of actin polymerization forces. 2016.

Table S1: Energy Minimization Parameters

| **Parameter** | **Setting** |
|---|---|
| Algorithm | Gradient descent |
| Energy tolerance | 10 kJ/mol/nm |
| Energy step size | 0.005nm |
| Number of steps | 15000 |
| Neighbor list update frequency | 1 step |
| Cutoff scheme | Verlet |
| Method to determine neighbor list | Grid |
| Short range force cut-off for neighbor list | 1.4nm |
| Electrostatics | Fast smooth Particle-Mesh Ewald (SPME) |
| Short-range electrostatic cut-off | 1.4nm |
| Short-range Van der Waals cut-off | 1.4nm |

Table S2: NVT Simulation Parameters

| **Parameter** | **Setting** |
|---|---|
| Time step | 2fs |
| Number of steps (Time) | 5000000 (10ns) |
| Integrator | Leapfrog algorithm |
| Constraint Algorithm | LINear Constraint Solver (LINCS) |
| Constraints | H-bonds constrained |
| Cutoff scheme | Verlet (Buffered neighbor searching) |
| Short-range electrostatic cutoff | 1.0nm |
| Short-range van der Waals cutoff | 1.0nm |
| Electrostatics | Fast smooth Particle-Mesh Ewald (SPME) |
| Interpolation order | Cubic |
| Grid spacing for fast Fourier Transform | 0.16nm |
| Temperature coupling | V-rescale (modified Berendsen themostat) |
| Reference temperature | 310K |
| Temperature time constant | 0.1ps |
| Temperature coupled groups | Protein and non-protein |
| Pressure coupling | Off |
| Dispersion correction | long range dispersion corrections for energy and pressure |
| Velocity generation | On |
| Temperature for velocity generation | 310K |

Table S3: NPT Simulation Parameters

| **Parameter** | **Setting** |
|---|---|
| Time step | 2fs |
| Number of steps (Time) | 5000000 (10ns) |
| Integrator | Leapfrog algorithm |
| Constraint Algorithm | LINear Constraint Solver (LINCS) |
| Constraints | H-bonds constrained |
| Cutoff scheme | Verlet (Buffered neighbor searching) |
| Short-range electrostatic cutoff | 1.0nm |
| Short-range van der Waals cutoff | 1.0nm |
| Electrostatics | Fast smooth Particle-Mesh Ewald (SPME) |
| Interpolation order | Cubic |
| Grid spacing for fast Fourier Transform | 0.16nm |
| Temperature coupling | V-rescale (modified Berendsen themostat) |
| Reference temperature | 310K |
| Temperature time constant | 0.1ps |
| Temperature coupled groups | Protein and non-protein |
| Pressure coupling | Isotropic Parrinello-Rahman |
| Pressure time constant | 1.0ps |
| Reference pressure | 1.0bar |
| Compressibility | 4.5e-5 bar-1 |
| Dispersion correction | long range dispersion corrections for energy and pressure |
| Velocity generation | Off |

Table S4: Steered Molecular Dynamics Parameters

| **Parameter** | **Setting** |
|---|---|
| Time step | 2fs |
| Number of steps (Time) for 1nm/ns | 12500000 (25ns) |
| Number of steps (Time) for 10nm/ns | 1500000 (3ns) |
| Integrator | Leapfrog algorithm |
| Constraint Algorithm | LINear Constraint Solver (LINCS) |
| Constraints | H-bonds constrained |
| Cutoff scheme | Verlet (Buffered neighbor searching) |
| Short-range neighbor list cutoff | 1.4nm |
| Short-range electrostatic cutoff | 1.4nm |
| Short-range van der Waals cutoff | 1.4nm |
| Electrostatics | Fast smooth Particle-Mesh Ewald (SPME) |
| Interpolation order | Cubic |
| Grid spacing for fast Fourier Transform | 0.12nm |
| Temperature coupling | Nosé-Hoover |
| Reference temperature | 310K |
| Temperature time constant | 1.0ps |
| Temperature coupled groups | Protein and non-protein |
| Pressure coupling | Off |
| Dispersion correction | long range dispersion corrections for energy and pressure |
| Velocity generation | Off |
| Harmonic potential | Umbrella |
| Force constant | 50 kJ/mol-nm$^2$ |
| Pull direction | y-direction (vertical) |
| Pull rate for 1nm/ns | 0.001nm/ps = 1nm/ns |
| Pull rate for 10nm/ns | 0.010nm/ps = 10nm/ns |

Table S5: Force Distribution Analysis Parameter Settings

| **Parameter** | **Setting** |
|---|---|
| Pairwise forces | Summed |
| Pairwise groups | Protein |
| Residue based calculation | Punctual Stress |
| Pairwise force type | Coulombic interactions only |

Table S6: Whole-cell Model Parameter Settings

| **Parameter** | **Variable** | **Setting** |
|---|---|---|
| Substrate modulus | $\mu_s$ | 1 MPa |
| Cell modulus | $\mu_c$ | 1 kPa |
| Max $\alpha_5\beta_1$-FN concentration | $\rho_{i,max}$ | $100\mu\text{m}^{-2}$ |
| Time step | $\delta t$ | 0.0005s |
| Catch-slip bond parameters | $K_a$ | 0.004 $s^{-1}$ |
| | $K_b$ | $10s^{-1}$ |
| | $F_a$ | $15pN$ |
| | $F_b$ | $15pN$ |

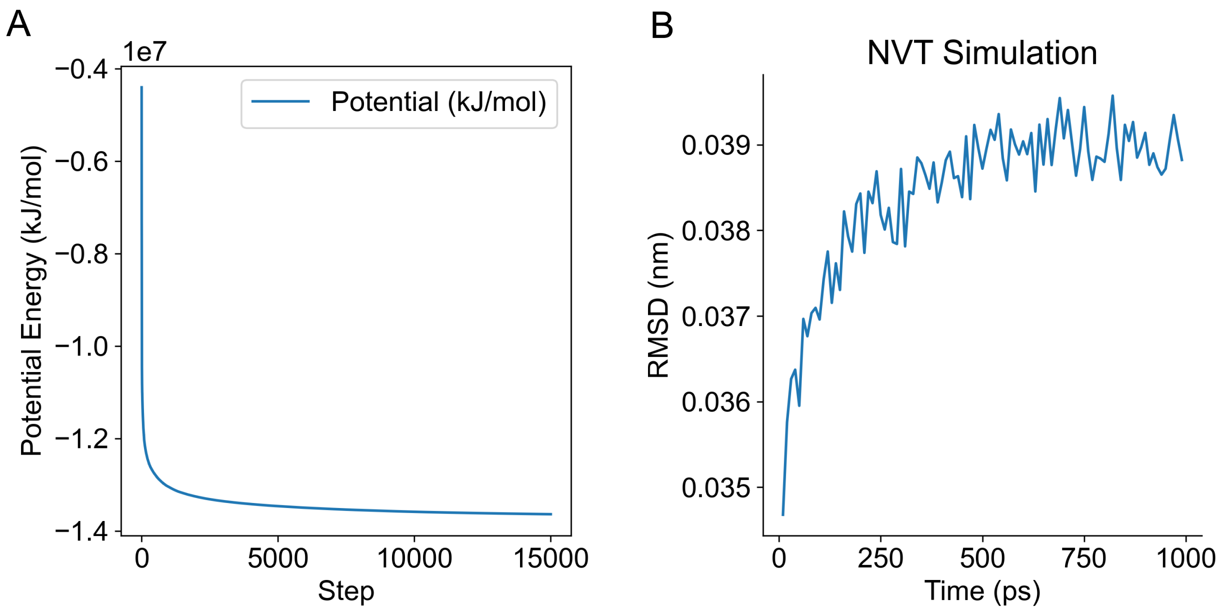


Figure S1: A) Energy minimization and B) root-mean-square deviation (RMSD) during 1ns NVT simulation of $\alpha_5\beta_1$-FN

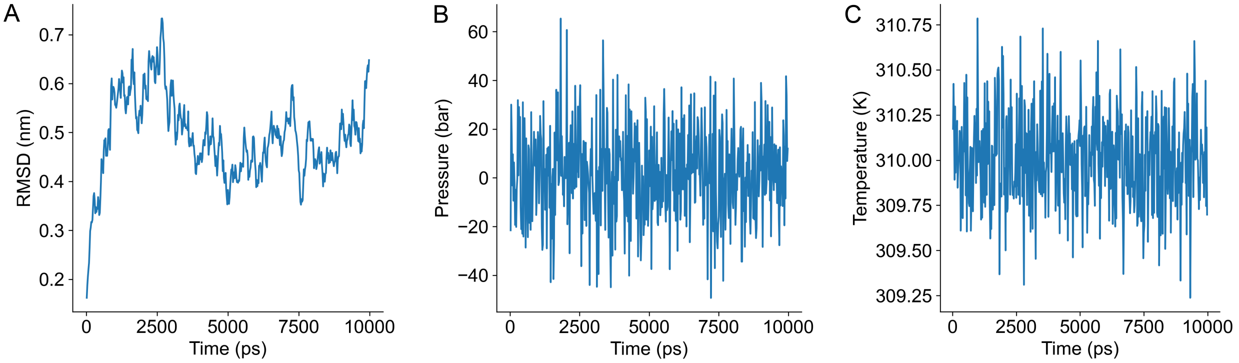


Figure S2: A) RMSD, B) Pressure, and C) Temperature of $\alpha_5\beta_1$-FN after 10ns NPT simulation indicative of an equilibrated system

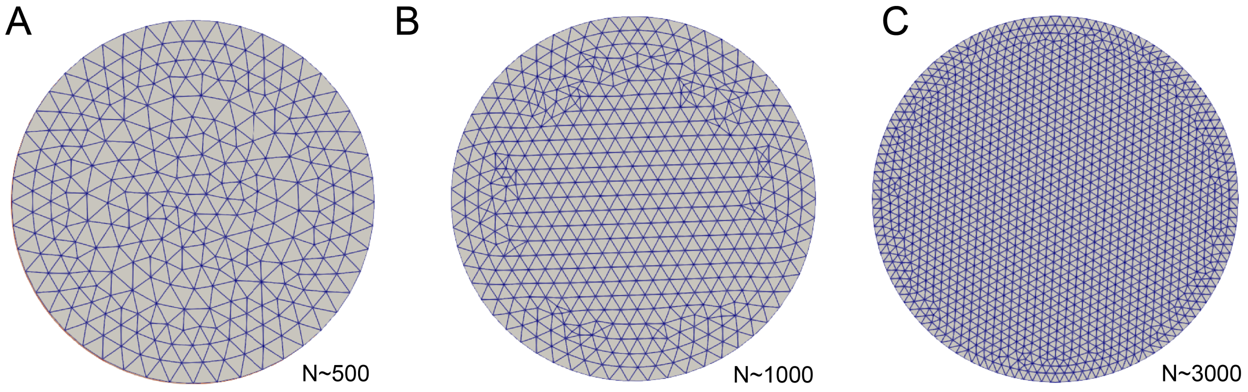


Figure S3: A) 500-element mesh. B) 1000-element mesh. C) 3000-element mesh.

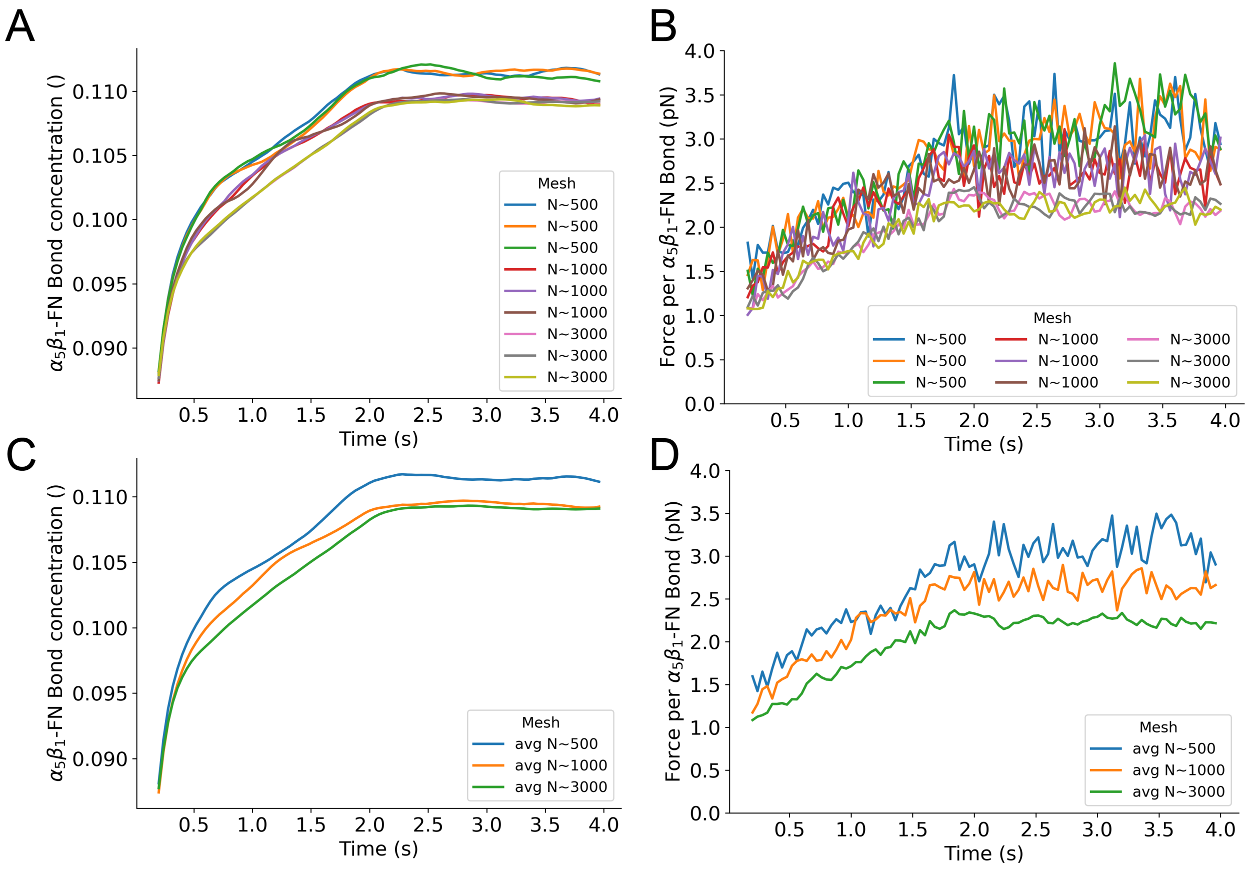


Figure S4: Mesh convergence (n=3 runs per mesh) of A) Bond concentration and B) force per bond. Average of C) bond concentration and D) force per bond across three runs. We chose the constant $k_{int}$=31pN/nm at a timestep of 100$\mu$s to run the convergence analysis.

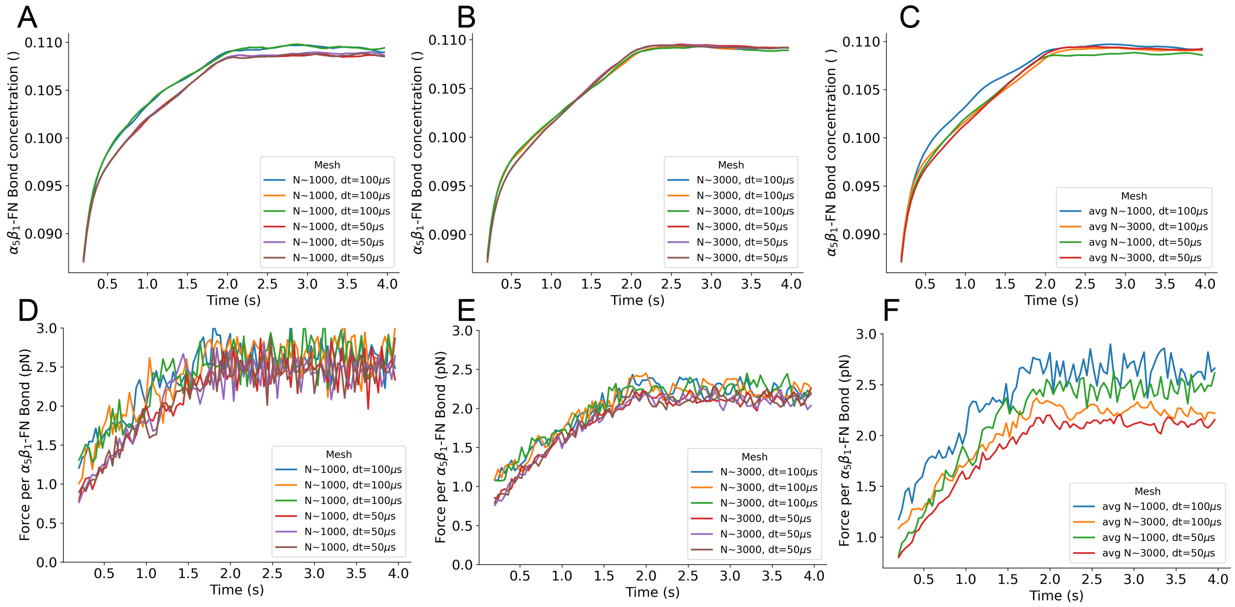


Figure S5: Timestep convergence (n=3 runs at 100$\mu$s and 50$\mu$s per 1000-element and 3000-element mesh). Bond concentration from A) three runs of 1000-element mesh at a 100$\mu$s and 50$\mu$s timestep. B) Three runs of 3000-element mesh at a 100$\mu$s and 50$\mu$s timestep. C) Comparison between time-averaged 1000-element and 3000-element mesh runs at a 100$\mu$s and 50$\mu$s timestep. Force per bond from D) three runs of 1000-element mesh at a 100$\mu$s and 50$\mu$s timestep. E) Three runs of 3000-element mesh at a 100$\mu$s and 50$\mu$s timestep. F) Comparison between time-averaged 1000-element and 3000-element mesh runs at a 100$\mu$s and 50$\mu$s timestep. We chose the constant $k_{int}$=31pN/nm to run the convergence analysis.
